# Camostat mesylate inhibits SARS-CoV-2 activation by TMPRSS2-related proteases and its metabolite GBPA exerts antiviral activity

**DOI:** 10.1101/2020.08.05.237651

**Authors:** Markus Hoffmann, Heike Hofmann-Winkler, Joan C. Smith, Nadine Krüger, Lambert K. Sørensen, Ole S. Søgaard, Jørgen Bo Hasselstrøm, Michael Winkler, Tim Hempel, Lluís Raich, Simon Olsson, Takashi Yamazoe, Katsura Yamatsuta, Hirotaka Mizuno, Stephan Ludwig, Frank Noé, Jason M. Sheltzer, Mads Kjolby, Stefan Pöhlmann

## Abstract

Antiviral therapy is urgently needed to combat the coronavirus disease 2019 (COVID-19) pandemic, which is caused by severe acute respiratory syndrome coronavirus 2 (SARS-CoV-2). The protease inhibitor camostat mesylate inhibits SARS-CoV-2 infection of lung cells by blocking the virus-activating host cell protease TMPRSS2. Camostat mesylate has been approved for treatment of pancreatitis in Japan and is currently being repurposed for COVID-19 treatment. However, potential mechanisms of viral resistance as well as camostat mesylate metabolization and antiviral activity of metabolites are unclear. Here, we show that SARS-CoV-2 can employ TMPRSS2-related host cell proteases for activation and that several of them are expressed in viral target cells. However, entry mediated by these proteases was blocked by camostat mesylate. The camostat metabolite GBPA inhibited the activity of recombinant TMPRSS2 with reduced efficiency as compared to camostat mesylate and was rapidly generated in the presence of serum. Importantly, the infection experiments in which camostat mesylate was identified as a SARS-CoV-2 inhibitor involved preincubation of target cells with camostat mesylate in the presence of serum for 2 h and thus allowed conversion of camostat mesylate into GBPA. Indeed, when the antiviral activities of GBPA and camostat mesylate were compared in this setting, no major differences were identified. Our results indicate that use of TMPRSS2-related proteases for entry into target cells will not render SARS-CoV-2 camostat mesylate resistant. Moreover, the present and previous findings suggest that the peak concentrations of GBPA established after the clinically approved camostat mesylate dose (600 mg/day) will result in antiviral activity.

## INTRODUCTION

The outbreak of the novel coronavirus severe acute respiratory syndrome coronavirus 2 (SARS-CoV-2) in the city of Wuhan, China, in the winter of 2019 and its subsequent pandemic spread has resulted in more than 14 million cases of coronavirus disease 2019 and more than 600.00 deaths (*1*). Antivirals designed to combat SARS-CoV-2 are not available and repurposing of existing drugs developed against other diseases is considered the fastest option to close this gap (*2*). Remdesivir, a drug generated to inhibit Ebola virus infection, has recently been shown to reduce the duration of hospitalization for COVID-19 (*3*). However, the drug failed to reduce fatality significantly (*3*) and beneficial effects were not observed in a previous clinical trial (*4*), indicating that additional therapeutic options are needed.

We previously showed that the SARS-CoV-2 spike protein (S) uses the host cell factors angiotensin-converting enzyme 2 (ACE2) and transmembrane protease serine 2 (TMPRSS2) for entry into target cells (*5*). TMPRSS2 is a cellular type II transmembrane serine protease (TTSP) expressed in human respiratory epithelium that cleaves and thereby activates the viral S protein. Activation is essential for viral infectivity and we found that the protease inhibitor camostat mesylate, which is known to block TMPRSS2 activity (*6*), inhibits SARS-CoV-2 infection of lung cells (*5*). Camostat mesylate has been approved for treatment of pancreatitis in Japan (*7-9*) and it is currently being investigated as a treatment of COVID-19 in several clinical trials in Denmark, Israel and USA (NCT04321096, NCT04353284, NCT04355052, NCT04374019).

The activity of TMPRSS2 is essential for SARS-CoV and MERS-CoV lung infection and disease development (*10, 11*). Whether TMPRSS2-independent pathways for S protein activation exist and contribute to viral spread outside the lung is not fully understood. The S proteins of SARS-CoV-2 and several other coronaviruses can be activated by the pH-dependent endosomal cysteine protease cathepsin L in certain cell lines (*5, 12-15*). However, this auxiliary S protein activation pathway is not operative in the lung, likely due to low cathepsin L expression (*16*). Whether this pathway contributes to the recently reported extrapulmonary spread of SARS-CoV-2 is unknown (*17*). Similarly, it is unclear whether TTSPs other than TMPRSS2 can promote extrapulmonary SARS-CoV-2 spread. Finally, camostat mesylate is rapidly hydrolyzed into the active metabolite 4-(4-guanidinobenzoyloxy)phenylacetic acid (GBPA) in patients (*18-20*) but it is unknown to what extend GBPA inhibits TMPRSS2 activity.

Here, we identify TTSPs other than TMPRSS2 that can be used by SARS-CoV-2 for S protein activation and demonstrate that they are inhibited by camostat mesylate. Moreover, we provide evidence that camostat mesylate is rapidly converted into GBPA in cell culture and that GBPA inhibits SARS-CoV-2 entry with almost identical efficiency as compared to camostat mesylate when cells are preincubated with these compounds.

## RESULTS

### Identification of novel SARS-CoV-2 S protein activating proteases

The TTSP family comprises several enzymes which have previously been shown to activate surface glycoproteins of coronaviruses and other viruses, at least upon directed expression (*21-23*). Therefore, we asked whether the S protein of SARS-CoV-2 (SARS-2-S) can employ TTSPs other than TMPRSS2 for its activation. For this, we expressed different TTSPs along with the SARS-CoV-2 receptor, ACE2, in the otherwise poorly susceptible BHK-21 cells, treated the cells with ammonium chloride, which blocks the cathepsin L-dependent, auxiliary activation pathway, and transduced the cells with previously described vesicular stomatitis virus (VSV)-based pseudotypes bearing SARS-2-S (*5*). Ammonium chloride treatment strongly reduced SARS-2-S-driven transduction and this effect was rescued upon expression of TMPRSS2 (Fig. 1), as expected. Notably, this effect was also efficiently rescued by expression of TMPRSS13 and, to a lesser degree, TMPRSS11D, TMPRSS11E and TMPRSS11F (Fig. 1). Thus, SARS-2-S can use diverse TTSPs for S protein activation upon overexpression, with S protein activation by TMPRSS13 being particularly robust.

**Fig. 1.**
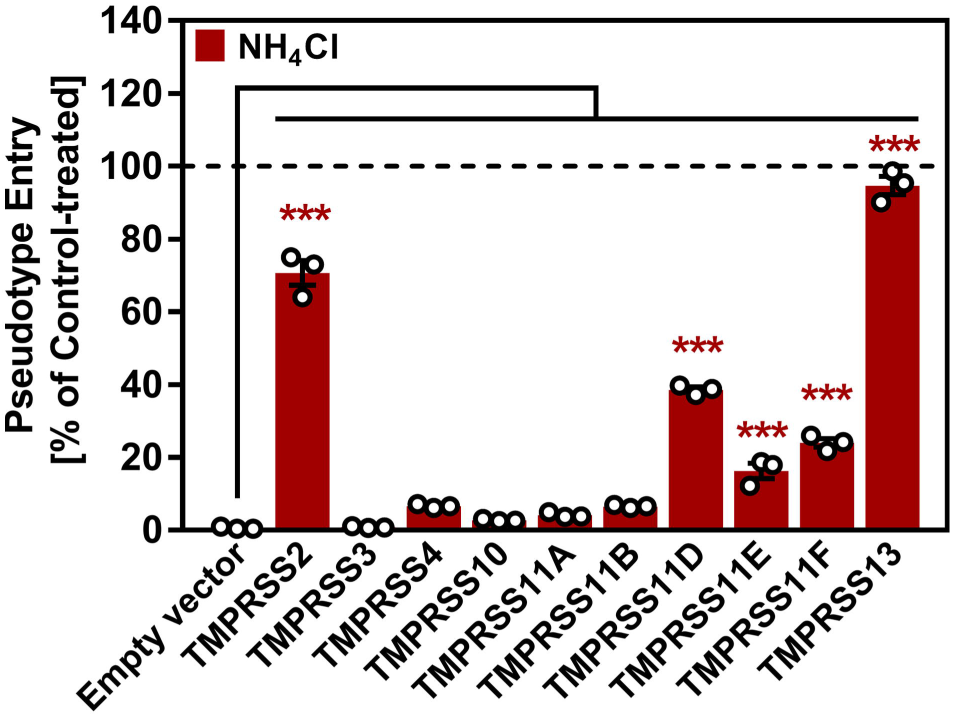
Different TTSPs can activate SARS-2-S in transfected cells. BHK-21 cells transiently expressing ACE2 and one of the indicated type-II transmembrane serine protease (or empty vector) were pre-incubated with either 50 mM ammonium chloride or DMSO (control, indicated by dashed line) for 2 h, before they were inoculated with pseudotype particles bearing SARS-2-S. At 16 h post inoculation, SARS-2-S-driven cell entry of viral pseudotypes was analyzed by measuring the activity of virus-encoded luciferase activity in cell lysates. Data were further normalized and entry efficiency in the absence of ammonium chloride was set as 100 %. Shown are the average (mean) data obtained from three biological replicates, each performed in quadruplicates. Error bars indicate the standard error of the mean (SEM). Statistical significance of differences in entry efficiency in the presence of ammonium chloride was analyzed by two-way analysis of variance (ANOVA) with Dunnett’s posttest.

### Several novel SARS-2-S activators are expressed in the airways and throat

In order to obtain insights into whether SARS-2-S activating TTSPs could contribute to viral spread in the infected host, we asked whether these enzymes are expressed in viral target cells. For this, we analyzed single-cell RNA-Seq datasets collected from human lungs (*24*) and airways (*25*). As previously reported (*26-31*), ACE2 was expressed in the lung epithelial compartment, particularly including alveolar type 2 cells, secretory (goblet/club) cells, and ciliated cells (Fig. 2A and Fig. S1). TMPRSS2 and TMPRSS13 were similarly expressed across epithelial cells, although TMPRSS13 expression was generally less robust. In contrast, expression of TMPRSS11-family members was only rarely detected (Fig. 2A). We found that 53% of ACE2^+^ cells in the lung co-express TMPRSS2, while 21% of ACE2^+^ cells do not express TMPRSS2 but do express another TTSP capable of activating SARS-CoV-2 (Fig. S1). Within the airways, we observed ACE2 expression in secretory cells, ciliated cells, and suprabasal cells in both the nasal turbinate and the trachea (Fig. 2B). Interestingly, the expression pattern of the TTSPs in the airways was largely distinct: TMPRSS2 was primarily expressed in ciliated and secretory cells, TMPRSS11D was primarily expressed in basal cells, TMPRSS11E was primarily expressed in ionocytes, and TMPRSS13 was primarily expressed in nasal secretory cells (Fig. 2B). Within this dataset, 21% of ACE2^+^ cells co-expressed TMPRSS2, while 24% of ACE2^+^ cells co-expressed a different TTSP (Fig. S1). In total, these results suggest that TMPRSS2 is the dominant SARS-CoV-2-activating protease in the lung, in keeping with findings made for SARS-CoV and MERS-CoV, while the virus may use other activating proteases for spread in the airways.

**Fig. 2.**
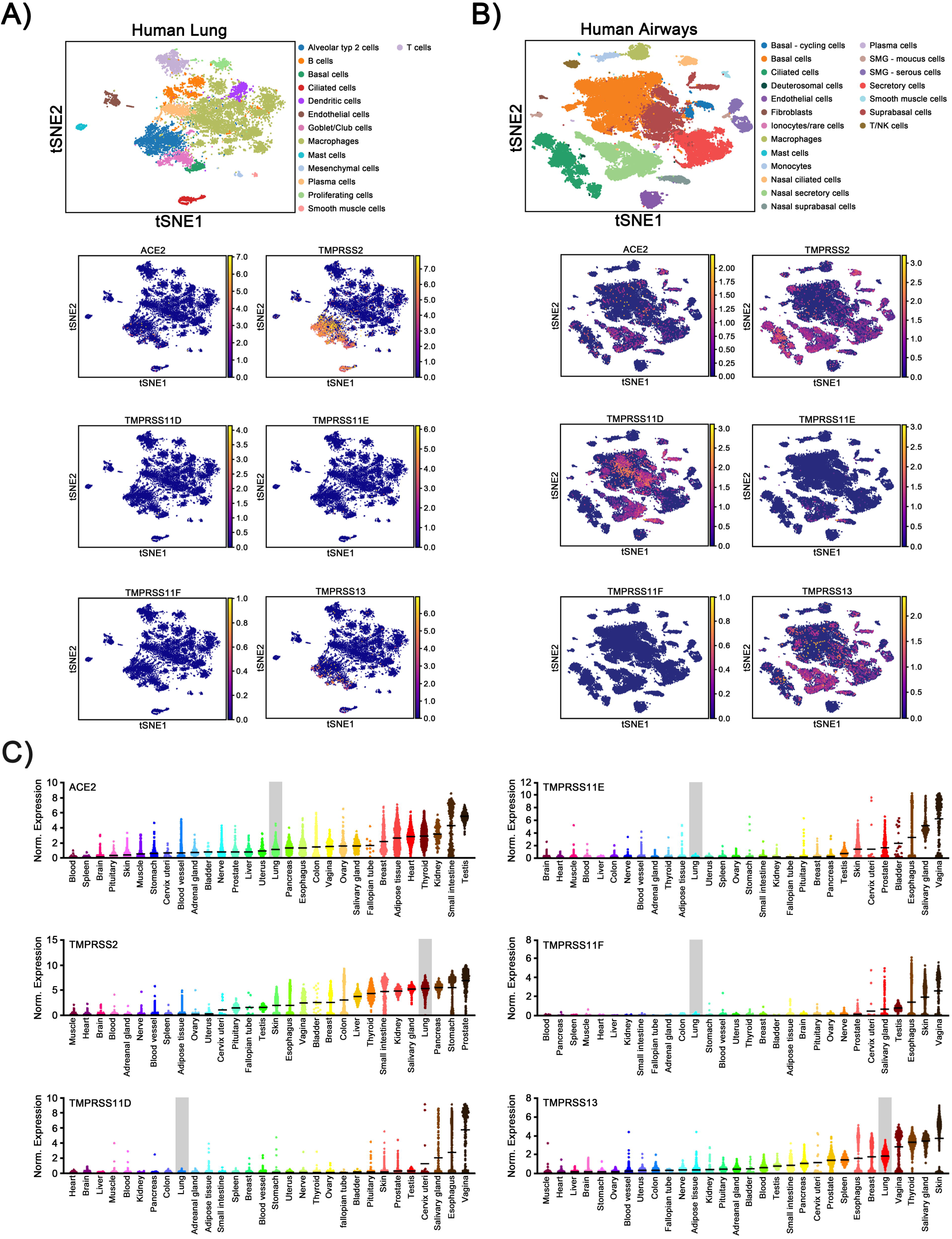
SARS-2-S activating proteases are expressed in lung and blood. (**A**) T-SNE clustering of cells from the human lung (*24*). Cells expressing the coronavirus receptor ACE2 are highlighted in the right panel. These panels are reproduced with permission from Smith et al. (*29*). Cells expressing various S-activating proteases in the human lung are highlighted. (**B**) T-SNE clustering of cells from the human airway (*25*). Cells expressing the coronavirus receptor ACE2 are highlighted in the right panel. Cells expressing various S-activating proteases in the human airway are highlighted. (**C**) Log2-normalized expression data of the indicated gene across different human tissues from the GTEx consortium (*33*).

A recent study provided evidence for extrapulmonary replication of SARS-CoV-2 in liver, colon, heart, kidney and blood in some patients (*17*). Therefore, we asked whether ACE2, TMPRSS2 and related SARS-2-S-activating proteases are expressed in these organs, using published resources (*32, 33*). Liver, colon, heart and kidney expressed robust levels of ACE2 (Fig. 2C). Similarly, TMPRSS2 expression in colon, liver and kidney was readily detectable, although expression levels were lower than those measured for lung (Fig. 2C). In contrast, little to no expression of TMPRSS11D, TMPRSS11E, TMPRSS11F, TMPRSS13 was detected in liver, colon, heart and kidney. Finally, TMPRSS13 was expressed in lung and blood cells and expression of TMPRSS11-family members was readily detectable in esophagus and salivary gland (Fig. 2C). Collectively, the TTSPs able to activate SARS-2-S were not expressed in appreciable levels in potential extrapulmonary targets of SARS-CoV-2. The only exceptions were TMPRS13 and TMPRSS11-family members that might contribute to SARS-CoV-2 infection of blood cells and to viral spread in the throat, respectively.

### Newly identified SARS-2-S activators are camostat mesylate sensitive

We next asked whether S protein activation by TTSP other than TMPRSS2 can be inhibited by camostat mesylate. To address this question, we performed the rescue assay as described above but investigated whether rescue can be blocked by camostat mesylate. In the absence of TTSP expression in target cells, ammonium chloride but not camostat mesylate reduced SARS-2-S-driven entry and the combination of both substances resulted in similar inhibition as observed upon ammonium chloride treatment alone (Fig. 3). These results are in agreement with only the cathepsin L-dependent auxiliary pathway being operative in control BHK-21 cells, in agreement with our published results (*5*). In TMPRSS2 transfected cells ammonium chloride did not efficiently block entry (Fig. 3), since under those conditions TMPRSS2 is available for S protein activation. Similarly, no entry inhibition was observed upon blockade of TMPRSS2 activity by camostat mesylate (Fig. 3), since the cathepsin L dependent activation pathway remained operative. Finally, the combination of ammonium chloride and camostat mesylate blocked entry into these cells (Fig. 3), in keeping with both activation pathways (cathepsin L and TMPRSS2) not being available under these conditions. Importantly, a comparable inhibition pattern was observed for all TTSPs able to activate SARS-2-S (Fig. 3), demonstrating that camostat mesylate will likely suppress SARS-CoV-2 activation by TMPRSS2 and TMPRSS2-related S protein activating serine proteases.

**Fig. 3.**
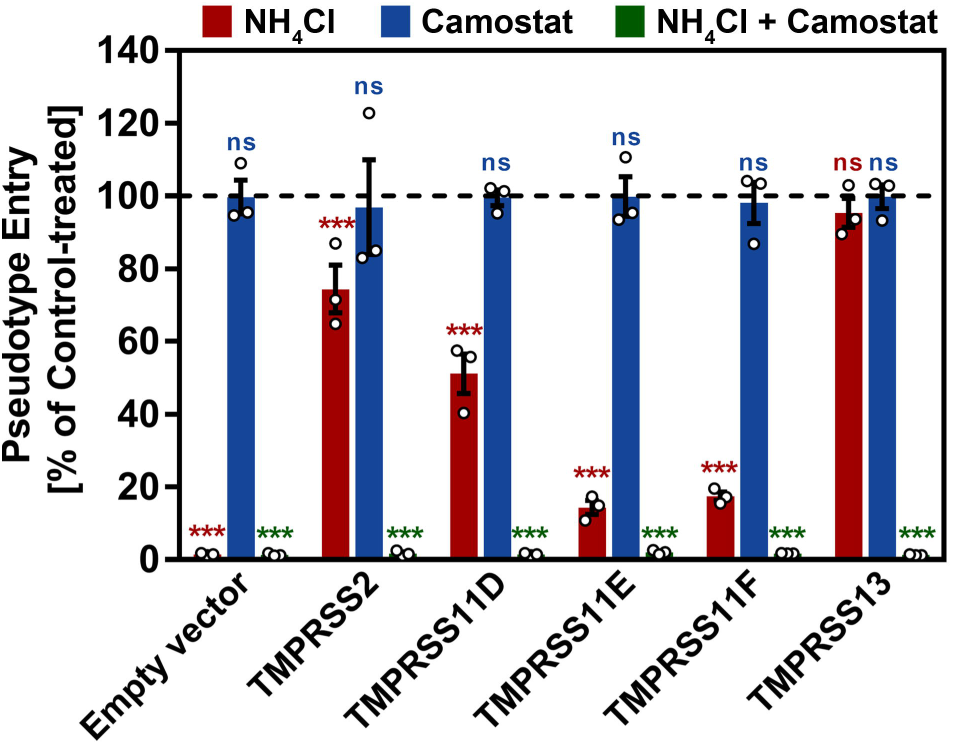
Activation of SARS-2-S by TMPRSS2-related proteases can be suppressed by camostat mesylate. The experiment was performed as described for figure 1 with the modifications that only TMPRSS2, TMPRSS11D, TMPRSS11E, TMPRSS11F and TMPRSS13 were investigated and target cells were pre-treated with either 50 mM ammonium chloride (red), 100 µM camostat mesylate (blue) or a combination of both (green). DMSO-treated cells served as controls. At 16 h post inoculation with viral particles bearing SARS-2-S, pseudotype entry was analyzed by measuring the activity of virus-encoded luciferase activity in cell lysates. Data were further normalized and entry efficiency into control-treated cells was set as 100 %. Shown are the average (mean) data obtained from three biological replicates, each performed in quadruplicates. Error bars indicate the SEM. Statistical significance of differences in entry efficiency in ammonium chloride-, camostat mesylate- or ammonium chloride + camostat mesylate-treated cells versus control-treated cells was analyzed by two-way ANOVA with Dunnett’s posttest (p > 0.5, not significant [ns], p ≤ 0.5, *; p ≤ 0.1, **; p ≤ 0.01, ***).

### The camostat mesylate metabolite GBPA shows reduced inhibition of recombinant TMPRSS2

Multiple studies show that camostat mesylate is rapidly converted into its active metabolite, 4-(4-guanidinobenzoyloxy)phenylacetic acid (GBPA) in animals and humans, followed by further conversion of GBPA into the inactive metabolite 4-guanidinobenzoic acid (GBA) (*18-20, 34*) (Fig. 6A). However, the capacity of GBPA to inhibit the enzymatic activity of TMPRSS2 has not been examined. To address this question, we compared inhibition of recombinant TMPRSS2 by camostat mesylate, GBPA and GBA. For this, we used FOY-251, a methanesulfonate of GBPA. We found that FOY-251 exerted a 10-fold reduced capacity to inhibit TMPRSS2 as compared to camostat mesylate, although both compounds completely suppressed TMPRSS2 activity at 1 µM or higher (Fig. 4). In contrast, GBA was less active (Fig. 4). Thus, FOY-251 blocks TMPRSS2 activity but with reduced efficiency as compared to camostat mesylate.

**Fig. 4.**
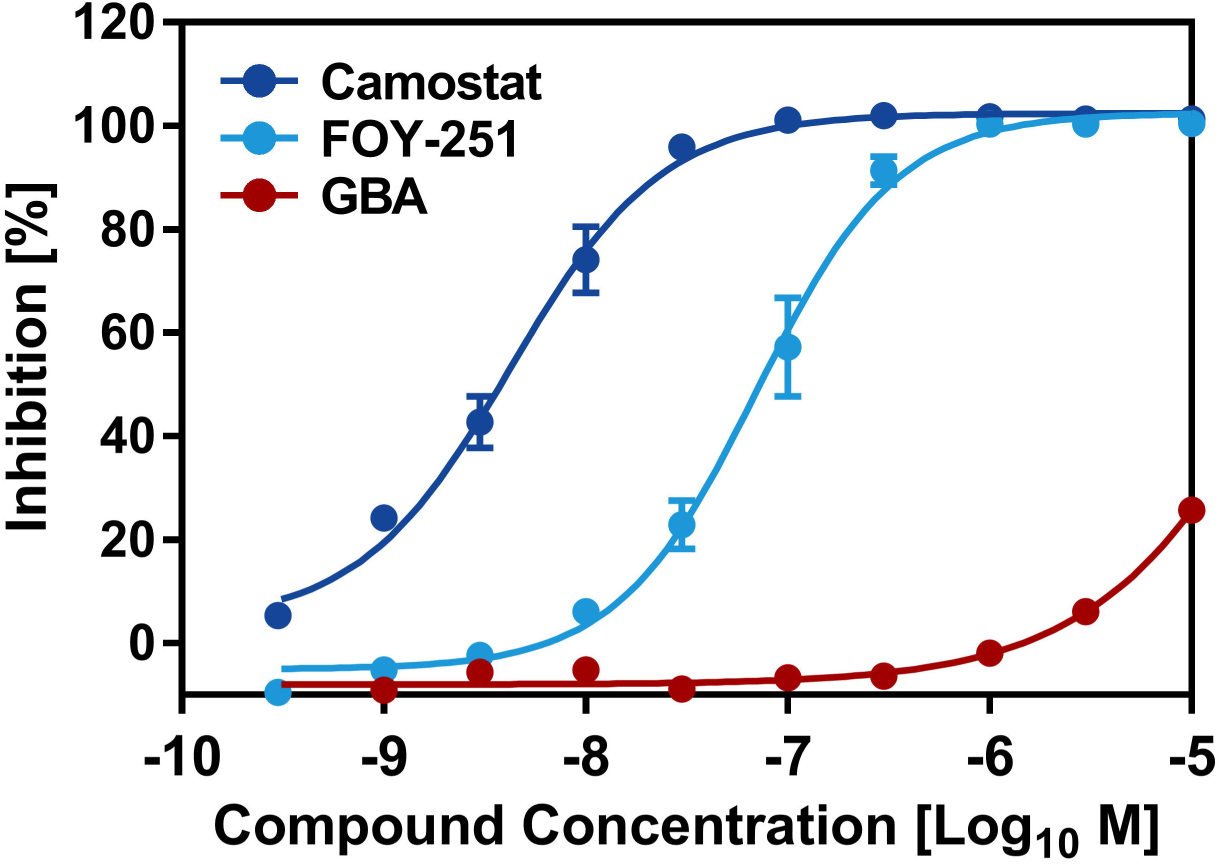
Camostat mesylate and FOY-251 inhibit the activity of recombinant TMPRSS2. TMPRSS2 cleaved Boc-Gln-Ala-Arg-MCA as substrate and produced the potent fluorophore, AMC(7-Amino-4-methylcoumarin). TMPRSS2 enzyme activity was evaluated by measuring the fluorescence intensity using Envision plate reader and all of the data were normalized against the intensity of the absence of test compounds. The concentration-response data for each test compound was plotted and modeled by a four-parameter logistic fit to determine the 50% inhibitory concentration (IC_50_) value. Inhibitory activity of camostat mesylate (blue), FOY-251(light blue) and GBA (red) against TMPRSS2 recombinant protein were visualized and curve fitting were performed using GraphPad Prism. The average of two independent experiments, each performed with quadruplicate (camostat mesylate and FOY-251) or duplicate samples (GBA) is shown. IC_50_ values were 4.2 nM (camostat mesylate), 70.3 nM (FOY-251), >10 µM (GBA).

**Fig. 5.**
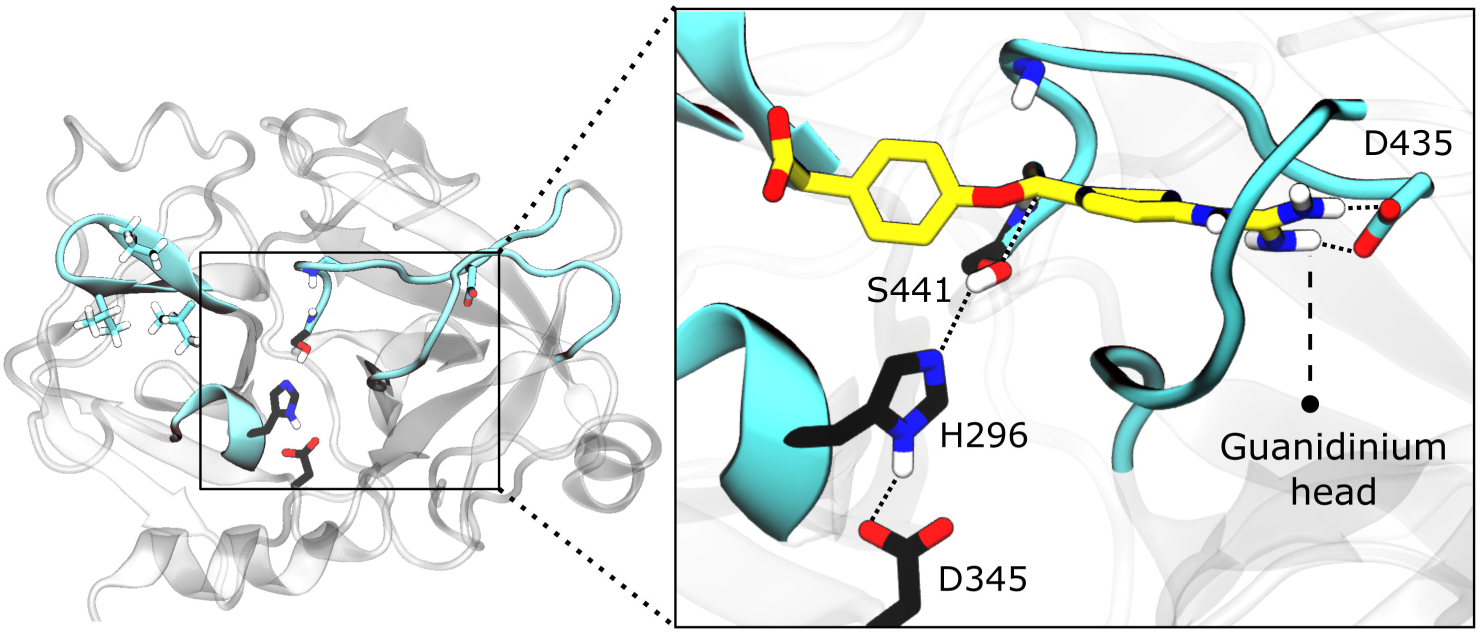
TMPRSS2 protease domain and GBPA interaction. A TMPRSS2 structure model is shown in the left panel, the active site is highlighted in cyan and catalytic triad residues are shown in black. The representative structure of GBPA bound to TMPRSS2 in a reactive complex is shown in the right panel. The GBPA guanidinium head forms a salt bridge with D435 inside the S1 pocket. This transient complex, which is similar for Camostat, is prone to be catalyzed at the ester bond interacting with Ser441, leading to a covalent complex with TMPRSS2 inhibited.

**Fig. 6.**
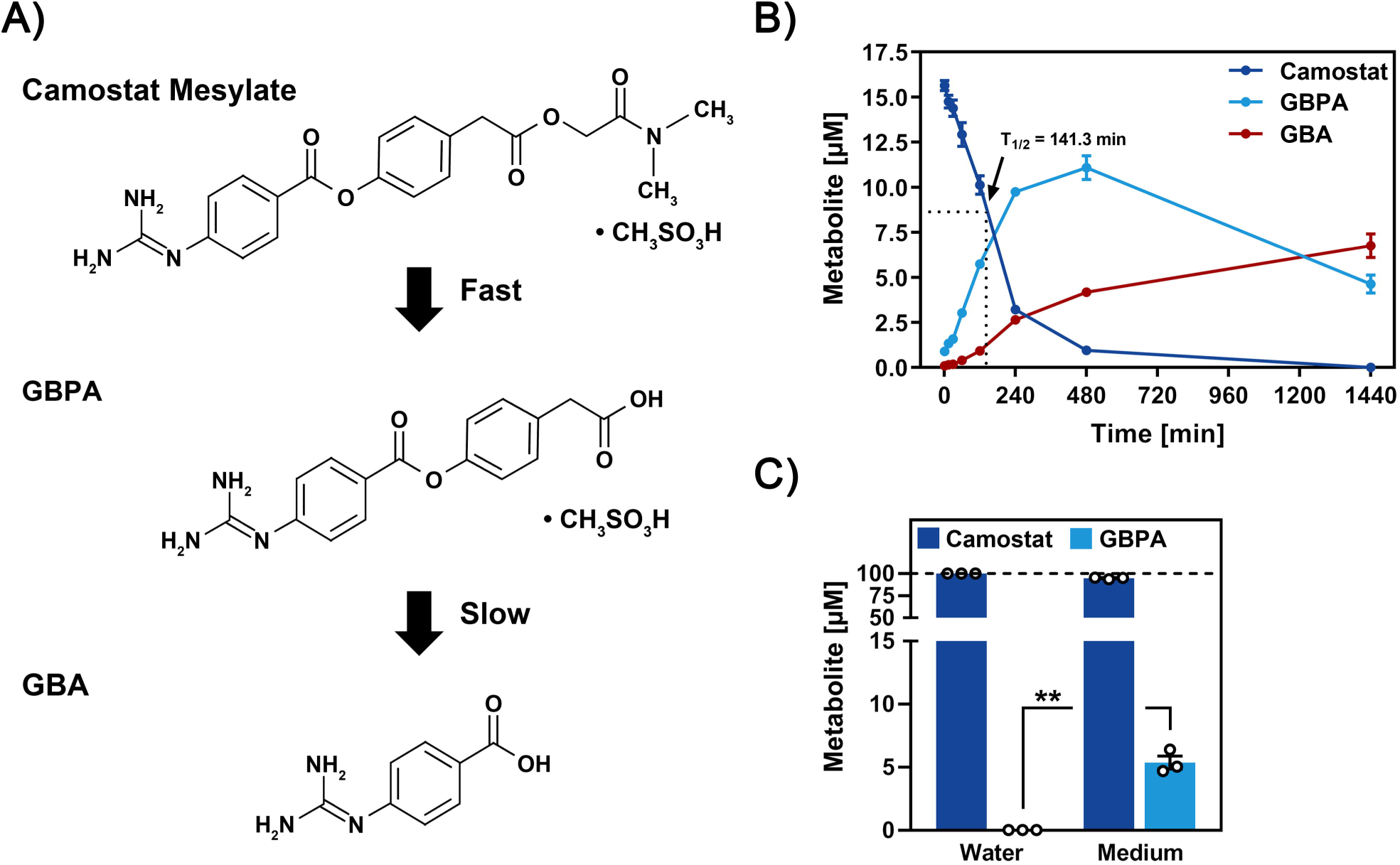
Camostat mesylate is rapidly converted into GBPA in the presence of cell culture medium. (**A**) Metabolization of camostat mesylate. (**B**) LC-MS/MS determination of camostat, GBPA and GBA in culture medium containing FCS. Camostat mesylate was added to FCS-containing culture medium at a concentration of 15 µM. Samples were taken after incubation for 1, 15, 30, 60, 120, 240, 480, and 1,440 min at 37 °C, snap-frozen and stored at −80 °C. Samples were analyzed by LC-MS/MS and quantified regarding their content of intact camostat mesylate and its metabolites GBPA (active) and GBA (inactive). Presented are the mean (average) data from three biological replicates (single samples). Error bars indicate the SEM. The turnover time that is required to cause metabolization of 50 % of camostat mesylate (T_1/2_) was further calculated by a non-linear regression model and was determined to be 141.3 min (95 % confidence interval = 116.5 to 171.7 min). (**C**) Relative levels of camostat mesylate and GBPA after incubation of 15 µM camostat mesylate in either water or FCS-containing culture medium. For normalization, the combined values of camostat mesylate and GBPA were set as 100 % and the relative fractions of the compounds were calculated. Presented are the mean (average) data from three biological replicates (single samples). Error bars indicate SEM. Statistical significance of differences in GBPA levels following incubation of camostat mesylate in either water or FCS-containing culture medium was analyzed by paired, two-tailed student’s t-test (p ≤ 0.01, **). Abbreviations: FOY-51/GBPA = 4-(4-guanidinobenzoyloxy)phenylacetic acid; GBA = 4-guanidinobenzoic acid.

In order to obtain insights into the reduced inhibitor activity of FOY-251, we investigated TMPRSS2 inhibition by GBPA on the molecular level. For this, we used a combination of extensive all-atom molecular dynamics (MD) simulations and Markov modeling of the TMPRSS2-GBPA complex (*35*). Guanidinobenzoate-containing drugs such as camostat mesylate and GBPA inhibit TMPRSS2 by first forming a noncovalent precomplex which is then catalyzed to form a long-lived covalent complex that is the main source of inhibition (*36*). However, the population of the short-lived precomplex directly relates to the inhibitory activity (*35*). By computing the TMPRSS2-GBPA binding kinetics (*35*), we find that (i) the noncovalent TMPRSS2-GBPA complex is metastable, rendering it suitable to form a covalent inhibitory complex, and (ii) its population is 40% lower compared to camostat at equal drug concentrations, consistent with the finding that FOY-251 is a viable but less potent inhibitor (Fig. 4).

Structurally, we find that GBPA binds in the same manner as camostat (Fig. 5, (*35*)). The main stabilizing interaction is its Guanidinium group binding into TMPRSS2’s S1 pocket which is stabilized by a transient salt bridge with Asp 435. The GBPA ester group can interact with the catalytic Ser 441, making it prone for catalysis and formation of the catalytic complex. The slightly lower stability of the GBPA compared to the camostat mesylate-TMPRSS2 complex is consistent with GBPA’s shorter tail which has less possibilities to interact with the hydrophobic patch on the TMPRSS2 binding site shown in Fig. 5, left panel.

### Rapid conversion of camostat mesylate to GBPA in cell culture

Although camostat mesylate is rapidly metabolized in animals and humans, it is less clear whether conversion of camostat mesylate into GBPA and GBA also occurs in cell culture. We addressed this question by exposing camostat mesylate to culture medium containing fetal calf serum (FCS), which is standardly used for cell culture, followed by mass spectrometric quantification of camostat mesylate and GBPA levels. Camostat mesylate levels rapidly declined with a half-life of approximately 2 h and the compound being barely detectable after 8 h (Fig. 6B). Conversely, the levels of the camostat mesylate metabolite GBPA increased rapidly, with peak levels attained at 8 h, and then remained relatively stable (Fig. 6B). Finally, the rapid metabolization of camostat mesylate into GBPA in the presence of serum was further confirmed by incubation of camostat mesylate in either water or FCS-containing culture medium for 1 min followed by quantification of camostat mesylate and GBPA levels. While GBPA levels were at background level when camostat mesylate was incubated in water, ∼5.4 % of camostat mesylate was metabolized into GBPA when incubated in FCS-containing culture medium (Fig. 6C). Thus, camostat mesylate is rapidly converted into GBPA under standard cell culture conditions, but the conversion is slower than what is observed in humans (*20*).

### Camostat mesylate and FOY-251 inhibit SARS-CoV-2 infection with comparable efficiency

We finally compared the antiviral activity of camostat mesylate and FOY-251, the methanesulfonate of GBPA, in cell culture. The reduced ability of FOY-251 to block the enzymatic activity of recombinant TMPRSS2 as compared to camostat mesylate would suggest that the compound should also exert reduced antiviral activity. On the other hand, analysis of antiviral activity encompasses preincubation of target cells with camostat mesylate for 2 h in the presence of FCS, which allows conversion of camostat mesylate into GBPA, as demonstrated above. Indeed, titration experiments with VSV pseudotypes and Calu-3 lung cells as targets revealed that entry inhibition by FOY-251 was only slightly reduced as compared to camostat mesylate, with EC50 values of 107 nM (camostat mesylate) and 178 nM (FOY-251) (Fig. 7). Moreover, no marked differences in inhibition of infection of Calu-3 cells with authentic SARS-CoV-2 were observed (Fig. 8). Thus, under the conditions chosen camostat mesylate and GBPA exerted comparable antiviral activity, likely due to conversion of camostat mesylate into GBPA.

**Fig. 7.**
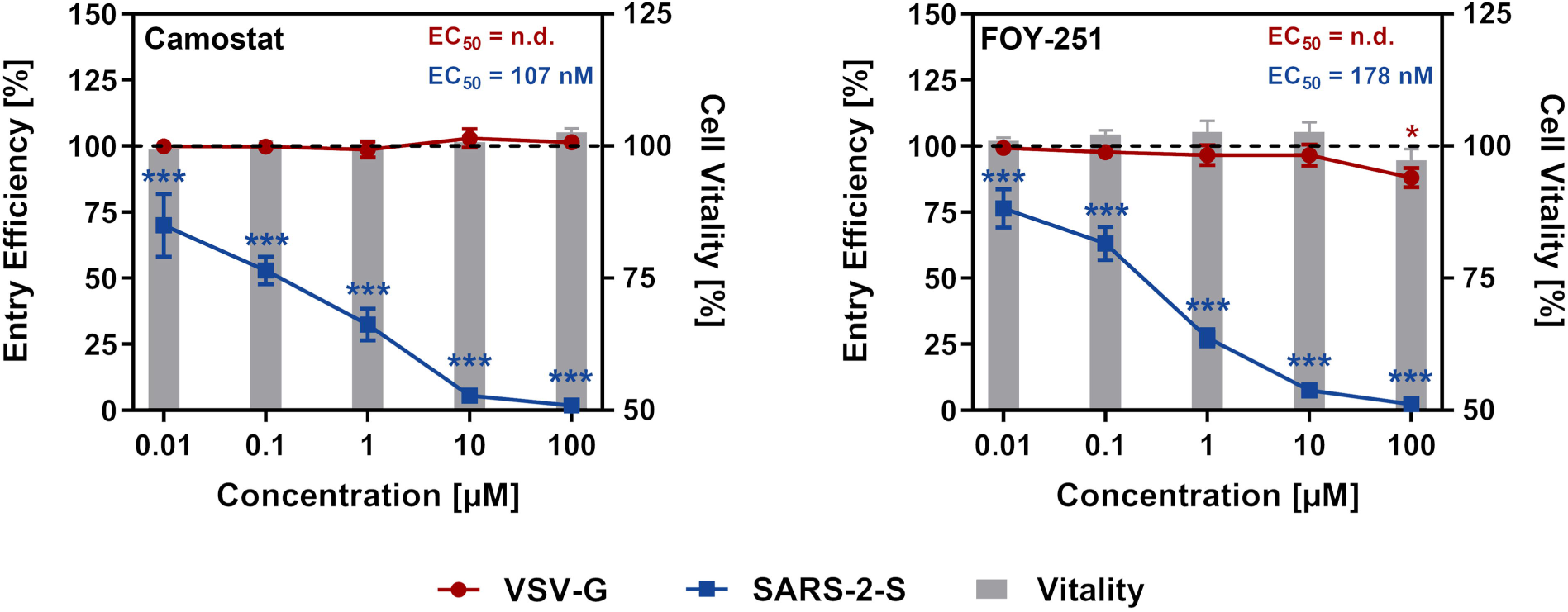
Camostat mesylate and FOY-251 inhibit SARS-2-S-driven cell entry with comparable efficiency. Calu-3 cells were pre-incubated with different concentrations of camostat mesylate (left panel), FOY-251 (right panel) or DMSO (control, indicated by dashed lines) for 2 h, before they were inoculated with pseudotype particles bearing VSV-G (red) or SARS-2-S (blue). Alternatively, in order to analyze potential negative effects of camostat mesylate and FOY-251 on cell vitality (grey bars), cells received medium instead of pseudotype particles and were further incubated. At 16 h post inoculation, pseudotype entry and cell vitality were analyzed by measuring the activity of virus-encoded luciferase activity in cell lysates or intracellular adenosine triphosphate levels (CellTiter-Glo assay), respectively. Data were further normalized against and entry efficiency/cell vitality in the absence of camostat mesylate and FOY-251 was set as 100 %. Shown are the average (mean) data obtained from three biological replicates, each performed in quadruplicates. Error bars indicate SEM. Statistical significance of differences in entry efficiency/cell vitality in camostat mesylate - or FOY-251-treated cells versus control-treated cells was analyzed by two-way ANOVA with Dunnett’s posttest (p > 0.5, not significant [ns], p ≤ 0.5, *; p ≤ 0.1, **; p ≤ 0.01, ***).

**Fig. 8.**
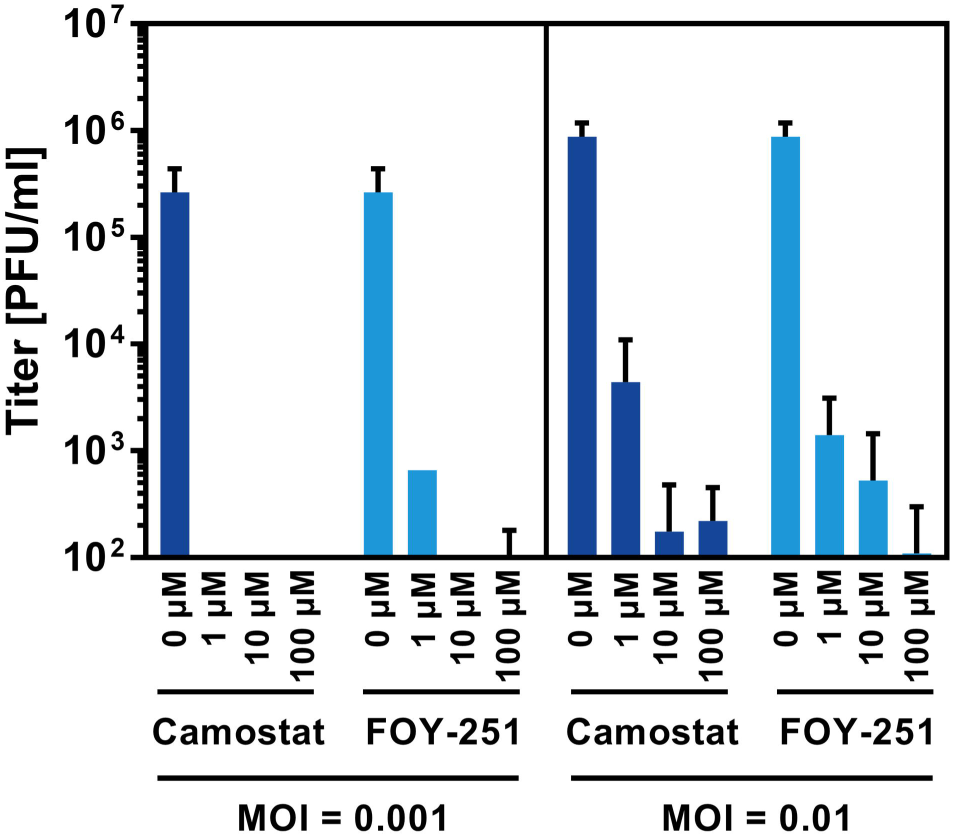
Camostat mesylate and FOY-251 inhibit SARS-CoV-2 infection with comparable efficiency. Calu-3 cells were pre-incubated for 2h with double concentration of camostat mesylate or FOY-251 as indicated. DMSO-treated cells served as control. Thereafter, cells were infected with SARS-CoV-2 at an MOI 0.001 or MOI 0.01 by adding the same volume of virus-containing culture medium to the inhibitor-treated cells. After 1 h of incubation, the inoculum was removed and cells were washed two times with PBS, before culture medium containing 1-fold inhibitor concentration was added. Culture supernatants were harvested at 24 h post infection, stored at −80°C and thereafter subjected to standard plaque formation assays using Vero-TMPRSS2 target cells and culture medium containing 1 % methyl cellulose. Plaques were counted at 48 h post infection and titers determined as plaque forming units per ml (pfu/ml). Presented are the data from a single experiment performed with technical triplicates and the results were confirmed in a separate experiment with another SARS-CoV-2 isolate. Error bars indicate the standard deviation.

## DISCUSSION

With the exception of remdesivir, which reduces disease duration (*3*), and dexamethasone, which reduces mortality in ICU patients by targeting inflammation (*37*), there are currently no drugs against COVID-19 with efficacy proven in clinical trials. We previously reported that the protease inhibitor camostat mesylate inhibits SARS-CoV-2 infection of cultured lung cells by blocking the virus-activating cellular protease TMPRSS2 (*5*). Camostat mesylate has been approved for human use in Japan and may thus constitute a COVID-19 treatment option. Here, we provide evidence that the virus can use TMPRSS2-related proteases for S protein activation and that these enzymes are also blocked by camostat mesylate. Moreover, we demonstrate that the camostat mesylate metabolite GBPA exhibits reduced ability to block enzymatic activity of purified, recombinant TMPRSS2 and is rapidly produced under cell culture conditions. The rapid conversion of camostat mesylate into GBPA likely accounts for our finding that both compounds exerted similar antiviral activity.

Knock-out of TMPRSS2 in mice markedly reduces SARS-CoV and MERS-CoV infection (*10*) and disease development, and similar findings have been reported for influenza A viruses (IAV) (*38-40*), which also use TMPRSS2 for glycoprotein activation (*41*). Thus, TMPRSS2 activity is essential for CoV and IAV infection of the lung. In contrast, several members of the TTSP family other than TMPRSS2 can activate CoV and IAV glycoproteins and support viral spread in cell culture, at least upon directed expression (*21, 23, 41*). Whether these TTSPs play a role in viral spread in the host is incompletely understood. For IAV, infection by H3N2 viruses were found not to be fully TMPRSS2 dependent (*38, 42*) and an auxiliary role of TMPRSS4 in spread and pathogenesis of H3N2 viruses has been reported (*43, 44*). Moreover, influenza B viruses can use a broad range of TTSPs in cell culture (*44, 45*) and a prominent role of TMPRSS2 in viral spread in type II pneumocytes has been reported (*46*) but viral spread in mice is TMPRSS2 independent (*44, 47*).

The present study shows that also SARS-CoV-2 can use TTSPs other than TMPRSS2 for S protein activation. Whether the TTSPs found here to activate SARS-2-S upon directed expression play a role in viral spread in the host remains to be investigated. Expression analyses suggest that they may. TMPRSS13 activated SARS-2-S with similar efficiency as TMPRSS2 and TMPRSS13 mRNA was found to be coexpressed with ACE2 in type II pneumocytes, goblet and club cells and basal cells. Moreover, TMPRSS13 was expressed in blood cells, which may constitute a target for SARS-CoV-2 infection in some patients. Finally, and most notably, SARS-S-2 activating TTSPs showed distinct expression patterns in the upper respiratory tract and several potential target cells coexpressed ACE2 jointly with a novel S protein activating TTSP but not TMPRSS2. Although viral spread supported by TMPRSS13 and potentially other SARS-2-S activating TTSPs could contribute to transmission and pathogenesis, it would still be sensitive to blockade by camostat mesylate. Thus, usage of auxiliary TTSPs for S protein activation would not confer camostat mesylate resistance to SARS-CoV-2.

In animal and humans camostat mesylate is rapidly hydrolyzed into the active metabolite 4-(4-guanidinobenzoyloxy) phenylacetic acid (GBPA), which is further hydrolyzed to 4-guanidinobenzoic acid (GBA) (*18-20*). GBPA was known to retain protease inhibitor activity (*34, 48*) but it was unclear whether GBPA would block TMPRSS2 activity with the same efficiency as camostat mesylate. Inhibition studies carried out with recombinant TMPRSS2 demonstrated that although GBPA robustly blocked TMPRSS2 activity, the compound was about 10-fold less active than camostat mesylate, which roughly matches results reported for other proteases (*49*). This finding raised the question whether camostat mesylate conversion into GBPA also occurs in cell culture systems used to assess antiviral activity of camostat mesylate. Indeed, camostat mesylate was rapidly converted into GBPA in the presence of serum which may account for camostat mesylate and FOY-251 exerting roughly comparable antiviral activity when cells were preincubated with these compounds for 2 h in the presence of serum. This has important implications for COVID-19 treatment, considering that continuous IV infusion of camostat mesylate (40 mg) resulted in a maximal plasma GBPA concentration of 0.22 µM and peak plasma concentrations of GBPA in humans upon oral intake of 200 mg camostat mesylate can reach 0.25 µM (http://www.shijiebiaopin.net/upload/product/201272318373223.PDF). Provided that concentrations in plasma and in respiratory epithelium are comparable, this would suggest that GBPA peak levels attained with the dosage approved for pancreatitis treatment (200 mg camostat mesylate three times a day) would be sufficient to exert antiviral activity.

Collectively, our results indicate that camostat mesylate constitutes a viable treatment option for COVID-19. Independent of its antiviral activity, camostat mesylate might reduce the uncontrolled cytokine release observed in severe COVID-19, since TMPRSS2 expression is required for robust cytokine release upon exposure of mice to polyIC (*10*).

## MATERIALS AND METHODS

### Cell culture

BHK-21 (baby hamster kidney; ATCC no. CCL-10) and HEK-293T (human embryonic kidney; DSMZ no. ACC 635) cells were cultivated in Dulbecco’s modified Eagle medium supplemented with 10 % fetal bovine serum (FCS, Biochrom), 100 U/mL of penicillin and 0.1 mg/mL of streptomycin (PAN-Biotech). Calu-3 cells (human lung adenocarcinoma) were cultivated in minimum essential medium (MEM) containing 10 % FCS (Biochrom), 100 U/mL of penicillin and 0.1 mg/mL of streptomycin (PAN-Biotech), 1x non-essential amino acid solution (from 100x stock, PAA) and 10mM sodium pyruvate (Thermo Fisher Scientific). Cell lines were incubated at 37 °C in a humidified atmosphere containing 5 % CO_2_. Transfection of 293T cells was performed by calcium-phosphate precipitation, while Lipofectamine LTX with Plus reagent (Thermo Fisher Scientific) was used for transfection of BHK-21 cells.

### Plasmids

We employed pCAGGS-based expression vectors for VSV-G, TMPRSS2, TMPRSS3, TMPRSS4, TMPRSS10, TMPRSS11A, TMPRSS11B, TMPRSS11D, TMPRSS11E, TMPRSS11F and TMPRSS13 that have either been previously described elsewhere or constructed on existing expression vectors (*21-23, 50-52*). All proteases contained an N-terminal cMYC-epitope tag. Further, we used pCG1-based expression vectors for human ACE2 (*53*) and a SARS-2-S variant with a truncated cytoplasmic tail for improved pseudotype particle production (deletion of last 18 amino acid residues,(*54*))

### Preparation of camostat mesylate and GBPA stocks

Camostat mesylate and GBPA were obtained from Ono pharmaceuticals Co., LTD. (Osaka/Japan) and reconstituted in DMSO to yield stock solutions of 100 mM. Stocks were stored at −20 °C, thawed immediately before the experiment and residual compound was discarded.

### Mass spectrometric quantification of camostat mesylate metabolization

Camostat mesylate was diluted to a concentration of ∼15 µM in either water or MEM containing 10 % FCS and incubated for 1 min (water and medium samples), 15 min, 30 min, 1 h, 2 h, 4 h, 8 h and 24 h (only medium samples) at 37 °C. Next, samples were snap-frozen and stored at −80 °C until camostat mesylate, GBPA and GBA levels were quantified by mass spectrometry. An ultra-high-performance liquid chromatography tandem mass spectrometry method using pneumatically assisted electrospray ionisation (UHPLC-ESI-MS/MS) was used for quantification of camostat and 4-(4-guanidinobenzoyloxy) phenylacetic acid (GBPA) in liquid samples. Calibrants based on blank sample were used for the construction of 8-point calibration curves. Calibrants were prepared with concentrations of 0.1, 1, 25, 50, 75, 100, 500 and 1000 µg/L of camostat and GBPA. In addition, a blank sample (a processed matrix sample without any added analyte) and a blank sample spiked with SIL-IS were included to verify the absence of detectable concentrations of the analytes. The calibration curves were created by weighted (1/x) regression analysis of the SIL-IS normalised peak areas (analyte area/IS area).

### Preparation of pseudotype particles

We employed a previously published protocol to generate vesicular stomatitis virus (VSV) pseudotype particles that is based on a replication-deficient VSV containing eGFP and firefly luciferase (FLuc) reporter genes, VSV*ΔG-FLuc (kindly provided by Gert Zimmer, Institute of Virology and Immunology IVI, Mittelhäusern/Switzerland) (*5, 55*). For this, HEK-293T cells were first transfected with expression vector for either SARS-2-S or VSV-G (or empty expression vector, control). At 24 h post transfection, cells were inoculated with VSV-G-transcomplemented VSV*ΔG-FLuc at a multiplicity of infection (MOI) of 3 and incubated for 1 h at 37 °C and 5 % CO_2_. Next, the inoculum was removed and cells were washed with PBS before fresh culture medium was added. In case of cells transfected with SARS-2-S-encoding vector or empty plasmid, the medium was spiked with anti-VSV-G antibody (supernatant of CRL-2700 cells, 1:1,000) in order to inactivate residual input virus containing VSV-G. At 16-18 h post inoculation, the culture supernatant was harvested and centrifuged (2,000 x g, 10 min) to remove cellular debris. Clarified supernatants containing pseudotype particles were aliquoted and stored at −80 °C until further use.

### Preparation of TMPRSS2 recombinant protein and substrate

Human TMPRSS2 (Recombinant N-terminus 6xHis, aa106-492) (Cat # LS-G57269-20) protein was acquired from LifeSpan Biosciences. Peptide Boc-Gln-Ala-Arg-MCA for the enzyme substrate was acquired from Peptide Institute, Inc.

### TMPRSS2 enzyme assay

All of different concentrations of test compounds were dissolved in DMSO and diluted with assay buffer (50 mM Tris-HCl pH 8.0, 154 mM NaCl) to the final DMSO concentration of 1%. Compound solution and Boc-Gln-Ala-Arg-MCA (10 μM final concentration) were added into the 384-well black plate (Greiner 784076). Then, enzyme reaction was started after adding TMPRSS2 recombinant protein to a final concentration of 2 μg/mL. Fluorescence intensity was read using the Envision plate reader with excitation: 380 nm and emission: 460 nm in 2 min intervals over 60 min at room temperature. The IC_50_ value was calculated based on the increasing rate of fluorescence intensity.

### Molecular dynamics simulations and Markov modeling

We used extensive all-atom molecular dynamics (MD) simulations of TMPRSS2 in complex with camostat as described in (*35*) starting from a homology model (*56*), in which drug binding and dissociation are sampled multiple times. We have then replaced camostat with GBPA and simulated a total of 50 µs MD with the same simulation setup as in [1] and used Markov modeling (*57*) to extract the dominant metastable binding modes of GBPA to the TMPRSS2 target on an atomistic scale. We estimate the binding kinetics of GBPA to the non-covalent complex by re-estimating the camostat Markov model described in (*35*) with the TMPRSS2-GBPA data. At the simulated drug concentration the association constant of GBPA is found to have a maximum likelihood estimate of 60% compared to that of camostat, resulting in a correspondingly lower inhibitory activity following the kinetic model of (*35*). Bootstrapping of trajectories under the constraint of comparable implied timescales yields a confidence interval of 51-100 % (68 % percentile).

### Transduction experiments

The day before transduction, BHK-21 cells were transfected with an expression vector for ACE2 and either empty expression plasmid (control) or expression vector encoding TMPRSS2, TMPRSS3, TMPRSS4, TMPRSS10, TMPRSS11A, TMPRSS11B, TMPRSS11D, TMPRSS11E, TMPRSS11F or TMPRSS13. For this, the old culture medium was removed and 50 µl/well of fresh culture medium were added. Next, transfection mixtures were prepared. For one well 0.1 µg of ACE2-encoding vector and 0.02 µg of protease-encoding vector (or empty plasmid) were mixed with 1 µl of Plus reagent, 50 µl of Opti-MEM medium (Thermo Fisher Scientific) and 1 µl of Lipofectamine LTX reagent. The transfection mix was vortexed and incubated for 30 min at room temperature before it was added to the cells. At 6 h post transfection, the transfection medium was replaced by fresh culture medium and the cells were further incubated for ∼18 h. Then, the cells were either pre-incubated for 2 h with inhibitor (50 mM ammonium chloride [Sigma-Aldrich], 100 µM camostat mesylate or a combination of both; cell treated with DMSO served as controls) before transduction or directly inoculated with pseudotype particles bearing SARS-2-S or cells. For transduction of Calu-3 cells, cells were pre-incubated for 2 h at 37 °C and 5 % CO_2_ with different concentrations (0.01, 0.1, 1, 10, 100 µM) of camostat mesylate, FOY-251 or DMSO (control), before they were inoculated with pseudotype particles bearing SARS-2-S or VSV-G. At 16 h post inoculation, transduction efficiency was analyzed by measuring the activity of virus-encoded FLuc in cell lysates. For this, the cell culture medium was removed and cells were incubated for 30 min with 1x concentrated Cell Culture Lysis Reagent (Promega), before cell lysates were transferred into white opaque-walled 96-well plates and luminescence was recorded (1 sec/sample) using a Hidex Sense plate luminometer (Hidex) and a commercial substrate (Beetle-Juice, PJK).

### Analysis of cell vitality

For the analysis of cell vitality of Calu-3 cells treated with camostat mesylate or FOY-251 the CellTiter-Glo Luminescent Cell Viability Assay kit (Promega) was used. For this, Calu-3 cells were grown in 96-well plates to reach ∼50% confluency, before they were incubated in the presence of different concentrations of camostat mesylate or FOY-251 for 24 h. Cells treated with DMSO (solvent control) served as controls. Following incubation, 100 µl of CellTiter-Glo substrate were added per well and the samples were incubated for 30 min on a rocking platform. In addition, fresh culture medium (without cells) was also incubated with CellTiter-Glo substrate in order to define the assay background. Following incubation, the samples were transferred into white opaque-walled 96-well plates and luminescence was recorded (200 msec/sample) using a Hidex Sense plate luminometer (Hidex).

### Infection of Calu-3 cells with authentic SARS-CoV-2

The SARS-CoV-2 isolate hCoV-19/Germany/FI1103201/2020 (GISAID accession EPI-ISL_463008) was isolated at the Institute of Virology, Muenster, Germany, from a patient returning from the Southern Tyrolean ski areas and propagated in Vero-TMPRSS2 cells. Calu-3 cells were pre-incubated for 2 h with 2-fold concentrated camostat mesylate or FOY-251 (2, 20 or 200 µM), or DMSO (control), before they were inoculated with SARS-CoV-2 at an MOI of 0.001 or 0.01. For this, the identical volume of virus-containing medium was added to the inhibitor-containing medium on the cells (resulting in 1-fold concentrated camostat mesylate or FOY-251; 1, 10 or 100 µM). Following 1 h of incubation at 37 °C and 5 % CO_2_, the culture supernatant was removed and cells were washed two times with excess PBS before culture medium containing 1-fold concentrated inhibitor was added. Supernatants were harvested at 24 h post inoculation and subjected to plaque titration. For this, confluent Vero-TMPRSS2 cells were inoculated with 10-fold serial dilutions of supernatant and incubated for 1 h 37 °C and 5 % CO_2_. Thereafter, the inoculum was removed and cells were incubated with culture medium containing 1 % (w/v) methyl cellulose. Plaques were counted at 48 h post infection and titers determined as plaque forming units per ml (pfu/ml).

### TTSP expression analysis

Bulk tissue expression data were obtained from the GTEx portal (*33*). Single-cell expression data from human lungs were obtained from GSE1229603. Only IPF and cryobiopsy lung explants were used in this analysis. Single-cell expression data from human airways was obtained from https://www.genomique.eu/cellbrowser/HCA/4. The single-cell data was analyzed as described in Smith et al (*29*). In short, dimensionality reduction and clustering were performed on normalized expression data in python using Scanpy and the Multicore-TSNE package (*58, 59*). Low quality cells were filtered out by removing cells with fewer than 500 detected genes. Highly variable genes were computed using the Seurat approach in Scanpy, and then used to calculate the principle component analysis. T-SNE and Leiden clustering were calculated using nearest neighbors, with parameters as described in the associated code. Cell clusters were labeled manually by comparing the expression patterns of established marker genes with the lists of differentially-expressed genes produced by Scanpy (*60-63*). The code used for performing these analyses is available at https://github.com/joan-smith/covid19-proteases/.

### Statistical analyses

All statistical analyses were performed using GraphPad Prism (version 8.4.2, GraphPad Software, Inc.). Statistical significance of differences between two datasets was analyzed by paired, two-tailed student’s t-test, while two-way analysis of variance (ANOVA) with Dunnett’s posttest was used for comparison of multiple datasets (the exact method used is stated in the figure legends). For the calculation of the turnover time required for metabolization of 50 % of camostat mesylate (T_1/2_) as well as the effective concentration 50 (EC50) values, which indicate the inhibitor concentration leading to 50 % reduction of transduction, non-linear fit regression models were used.

## SUPPLEMENTERY MATERIALS

Fig. S1, panel A. A track plot displaying the expression of ACE2, S-activating proteases, and several lineage-enriched genes in different lung cell populations obtained from Leiden clustering.

Fig. S1, panel B. A track plot displaying the expression of ACE2, S-activating proteases, and several lineage-enriched genes in different airway cell populations obtained from Leiden clustering.

Fig. S1, panel C. The percent of cells in the lung that express the indicated single gene or pair of genes are displayed.

Fig. S1, panel D. The percent of cells in the airway that express the indicated single gene or pair of genes are displayed.

## Acknowledgments

We are grateful for in-depth discussions with Katarina Elez, Robin Winter, Tuan Le, Moritz Hoffmann (FU Berlin) and the members of the JEDI COVID-19 grand challenge.

## Funding

Research in the Sheltzer Lab was supported by NIH grants 1DP5OD021385 and R01CA237652-01, a Damon Runyon-Rachleff Innovation award, an American Cancer Society Research Scholar Grant, and a grant from the New York Community Trust. The Noé lab was supported by Deutsche Forschungsgemeinschaft DFG (SFB/TRR 186, Project A12), the European Commission (ERC CoG 772230 “ScaleCell”), the Berlin Mathematics center MATH+ (AA1-6) and the federal ministry of education and research BMBF (BIFOLD). The Pöhlmann lab was supported by BMBF (RAPID Consortium, 01KI1723D). The Kjolby lab was supported by the Lundbeck Foundation (M.K., O.S.) and the Novo Nordisk Foundation (M.K.).

## Author contributions

M.H., H.M., F.N., J.M.S., M.K. and S.P. designed the study. M.H., H.H.-W., J.C.S., N.K., LK.S., O.S.S., J.B.H., T.H., L.R., S.O., T.Y., K.Y., and J.M.S., performed research. M.H., J.C.S., H.M., T.H., F.N. J.S.M., M.K. and S.P. analyzed the data. M.W. and S.L. provided essential reagents. M.H. and S.P. wrote the manuscript. All authors revised the manuscript.

## Competing interests

J.C.S. is a co-founder of Meliora Therapeutics and is an employee of Google, Inc. This work was performed outside of her affiliation with Google and used no proprietary knowledge or materials from Google. J.M.S. has received consulting fees from Ono Pharmaceuticals, is a member of the Advisory Board of Tyra Biosciences, and is a co-founder of Meliora Therapeutics. As part of its mission the Deutsches Primatenzentrum (German Primate Center) performs services for the scientific community including services for pharmaceutical companies resulting in fees being paid to the German Primate Center.

## Data availability statement

All data associated with this study are shown in the paper or the Supplementary Materials. All of the data used in this manuscript to determine protease expression are described in Table S1 of (*29*) and the code used for performing these analyses is available at github.com/joan-smith/covid19.

