## Supplementary figures and images for "Camostat mesylate inhibits SARS-CoV-2 activation by TMPRSS2-related proteases and its metabolite GBPA exerts antiviral activity"

### Supplementary Figure 1

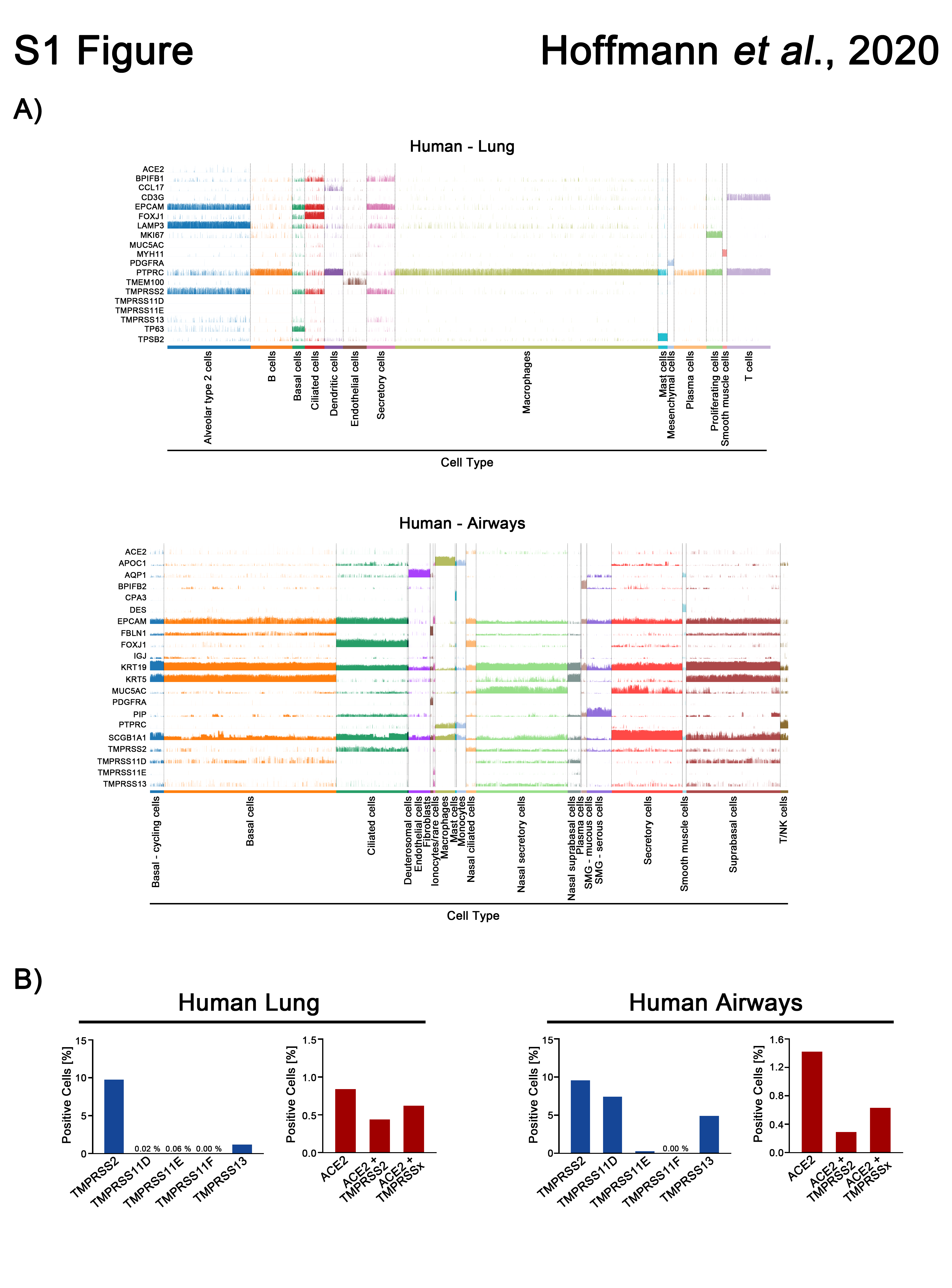
